# West Nile virus capsid protein promotes viral replication and pathogenesis through PKCα-dependent lamin phosphorylation and nuclear deformation

**DOI:** 10.64898/2026.07.17.739139

**Authors:** Keisuke Maezono, Passawat Thammahakin, Michiyo Kataoka, Tadaki Suzuki, Haruto Eguchi, Duong Thi Ngoc Thuy, Toshiro Yamaguchi, Akira Ota, Yukari Itakura, Koshiro Tabata, Hirofumi Sawa, Kentaro Yoshii, Hiroaki Kariwa, Shintaro Kobayashi

## Abstract

The genus *Orthoflavivirus* comprises several medically important pathogens such as the West Nile virus (WNV), which causes encephalitis in humans. Although viral replication occurs in the cytoplasm, the capsid (C) protein of the orthoflavivirus is localized to both the cytoplasm and nucleus. Nuclear C protein contributes to viral replication and disease progression. However, the underlying mechanisms remain unclear. Here, we investigated whether the WNV C protein induces nuclear deformation and examined the underlying mechanism. We also assessed the contribution of this deformation to viral replication and pathogenesis. WNV infection and C protein expression induced morphological alterations in the nuclear lamina, leading to nuclear deformation. C protein expression enhanced lamin phosphorylation and the disassembly of the polymerized lamin network. In addition, C protein interacted with protein kinase C alpha (PKCα) and localized PKCα near the nuclear lamina. Downregulation of PKCα expression inhibited C protein-induced lamin phosphorylation and nuclear deformation. In addition, both the downregulation of PKCα expression and pharmacological inhibition of PKC reduced WNV replication. In contrast, the expression of phosphorylation-deficient lamin mutants attenuated the inhibitory effect of downregulated PKCα expression on WNV replication. Furthermore, the pharmacological inhibition of PKC increased the survival rate of WNV-infected mice and suppressed both viral replication and nuclear deformation in the brain. Collectively, these results demonstrate that C protein remodels the nuclear lamina architecture through the PKCα−lamin pathway, and that virus-induced nuclear deformation contributes to WNV replication and pathogenesis.

**Author summary:** The West Nile virus (WNV), a neurotropic orthoflavivirus, causes severe neurological diseases in humans. In host cells, orthoflaviviruses exclusively replicate in the cytoplasm. However, their capsid (C) proteins are localized to both the nucleus and cytoplasm. Although the nuclear C protein has been implicated in viral replication and disease progression, its underlying mechanisms remain unclear. Here, we demonstrate that the WNV C protein induces nuclear deformation, accompanied by the phosphorylation of lamin and disassembly of the nuclear lamina, a structural scaffold that maintains the nuclear shape. The C protein promotes the localization of PKCα, a host kinase protein, near the nuclear lamina.

Suppression of PKCα expression or activity reduces lamin phosphorylation, nuclear deformation, and WNV replication. Importantly, pharmacological inhibition of PKC in WNV-infected mice reduced nuclear deformation and viral replication in the brain and improved survival rates. Collectively, our findings identify the host nucleus as an important site of WNV-host interaction and provide a new perspective that WNV, despite replicating in the cytoplasm, remodels host nuclear architecture to promote viral replication and pathogenesis.

## Introduction

The genus *Orthoflavivirus* in the family *Flaviviridae* comprises numerous arthropod-borne viruses of medical importance, including the West Nile virus (WNV), Japanese encephalitis virus (JEV), and tick-borne encephalitis virus (TBEV). In nature, WNV is maintained in an enzootic transmission cycle between mosquitoes and birds. In contrast, humans and horses are considered incidental dead-end hosts that become infected through mosquito bites [1]. WNV causes a wide range of clinical manifestations, including self-limiting febrile illness (West Nile fever) and severe neuroinvasive diseases such as meningoencephalitis and acute flaccid paralysis [2]. The number of reported cases of WNV infection has increased in recent years, particularly in Europe and the United States. This is consistent with the expanding geographical distribution of mosquito vectors [3]. Despite its increasing public health importance, no licensed vaccines or effective antiviral therapies are available for human WNV infections.

Orthoflaviviruses enter host cells *via* endocytosis and release their genomes into the cytoplasm [4]. The viral genome is an approximately 11 kb positive-sense, single-stranded RNA that encodes a single polyprotein. Viral and cellular proteases cleave this polyprotein into three structural proteins (C, prM/M, and E) and seven nonstructural proteins (NS1, NS2A, NS2B, NS3, NS4A, NS4B, and NS5) [5]. Viral genome replication, viral protein synthesis, and virion assembly occur in replication organelles derived from the endoplasmic reticulum membrane [5]. Following assembly, progeny viral particles are transported to the Golgi apparatus and released from the cell *via* the host secretory pathway [5,6]. Thus, the intracellular viral replication cycle of orthoflavivirus is thought to be completed within the cytoplasm. Nevertheless, the C protein of orthoflavivirus is localized not only in the cytoplasm but also in the nucleus [7–10]. Given the impairment of its nuclear localization attenuates viral replication and pathogenicity, C protein in the nucleus is considered important for the replication of orthoflavivirus [8,11]. However, the underlying molecular mechanisms remain unclear.

The nucleus houses the cellular genome and serves as a central hub for essential cellular activities, including DNA replication, transcription, and RNA processing [12]. The nucleus is spatially separated from the cytoplasm by the nuclear membrane. This nuclear membrane is composed of a double lipid bilayer, outer and inner nuclear membranes. In addition, it comprises nuclear pore complexes (NPCs) formed by approximately 30 distinct nucleoporins (Nups) [13]. NPC regulates the bidirectional transport of molecules across the nuclear membrane between the nucleus and cytoplasm [13]. The nuclear membrane is mechanically supported by the nuclear lamina, a meshwork formed by the polymerization of A- and B-type lamins and inner nuclear membrane proteins [14]. The A-type lamins—lamin A and lamin C (Lamin A/C)—are spliced isoforms encoded by *LMNA*. In contrast, B-type lamins (lamin B1 and lamin B2) are encoded by *LMNB1* and *LMNB2*, respectively [15,16].

Polymerization of the nuclear lamina is tightly regulated by the phosphorylation status of lamins. Multiple kinases regulate lamin phosphorylation under various cellular conditions. Host kinases such as protein kinase C (PKC) and cyclin-dependent kinase 1 mediate lamin phosphorylation during mitosis, leading to nuclear lamina disassembly and membrane remodeling [17–19]. Lamin phosphorylation can also be induced during interphase, including in response to viral infection [20–23]. For example, the UL97 protein of human cytomegalovirus localizes to the nuclear lamina and directly phosphorylates lamin [23].

Moreover, the ɣ134.5 protein of herpes simplex virus type 1, which alternates between the cytoplasm and nucleus, promotes lamin phosphorylation by interacting with PKC and redistributing it from the cytoplasm to the nucleus [21]. These observations suggest that nuclear-localized viral proteins modulate the integrity of the nuclear lamina.

Disruption of nuclear lamina integrity results in nuclear deformation characterized by blebbing, wrinkling, folding, and breakdown [24,25], which has been implicated in various human diseases, including Hutchinson-Gilford progeria syndrome, cardiomyopathy, and neurodegenerative diseases [26–30]. In addition to providing structural support, lamins interact with multiple binding partners, such as nups, inner nuclear membrane proteins, and chromatin. Thus, nuclear deformation likely contributes to the pathogenesis of various diseases by altering the cellular environment through impaired nucleocytoplasmic transport and/or alterations in gene expression [31–33]. However, whether nuclear deformation occurs during orthoflavivirus infection or contributes to viral replication remains unclear.

Based on these observations, we hypothesized that the WNV C protein alters nuclear morphology and facilitates viral replication. In this study, we investigated whether the WNV C protein induces nuclear deformation, examined the underlying mechanism, and assessed the contribution of this deformation to viral replication and pathogenesis. We demonstrate that the C protein induces nuclear deformation through its interaction with a host kinase, thereby promoting WNV replication and pathogenesis.

## Results

### WNV infection induced nuclear deformation

Morphological alternations in the nucleus have a critically influence on nuclear and cellular functions [25]. In WNV-infected cells, several viral proteins localize into the nucleus, and this nuclear localization contributes to efficient viral replication [7,34]. We therefore examined the morphology of the nucleus in WNV-infected cells by transmission electron microscopy (TEM). TEM analysis revealed marked alterations in the nuclear morphology of WNV-infected cells, with prominent deformation of the nuclear membrane characterized by discontinuities and bleb-like structures. In contrast, the nuclei in mock-infected cells displayed a smooth and continuous structure (Fig 1A). The number of WNV-infected cells with nuclear membrane deformation was higher (22.4%) than that of mock cells (0%) (Fig 1A).

**Fig 1.**
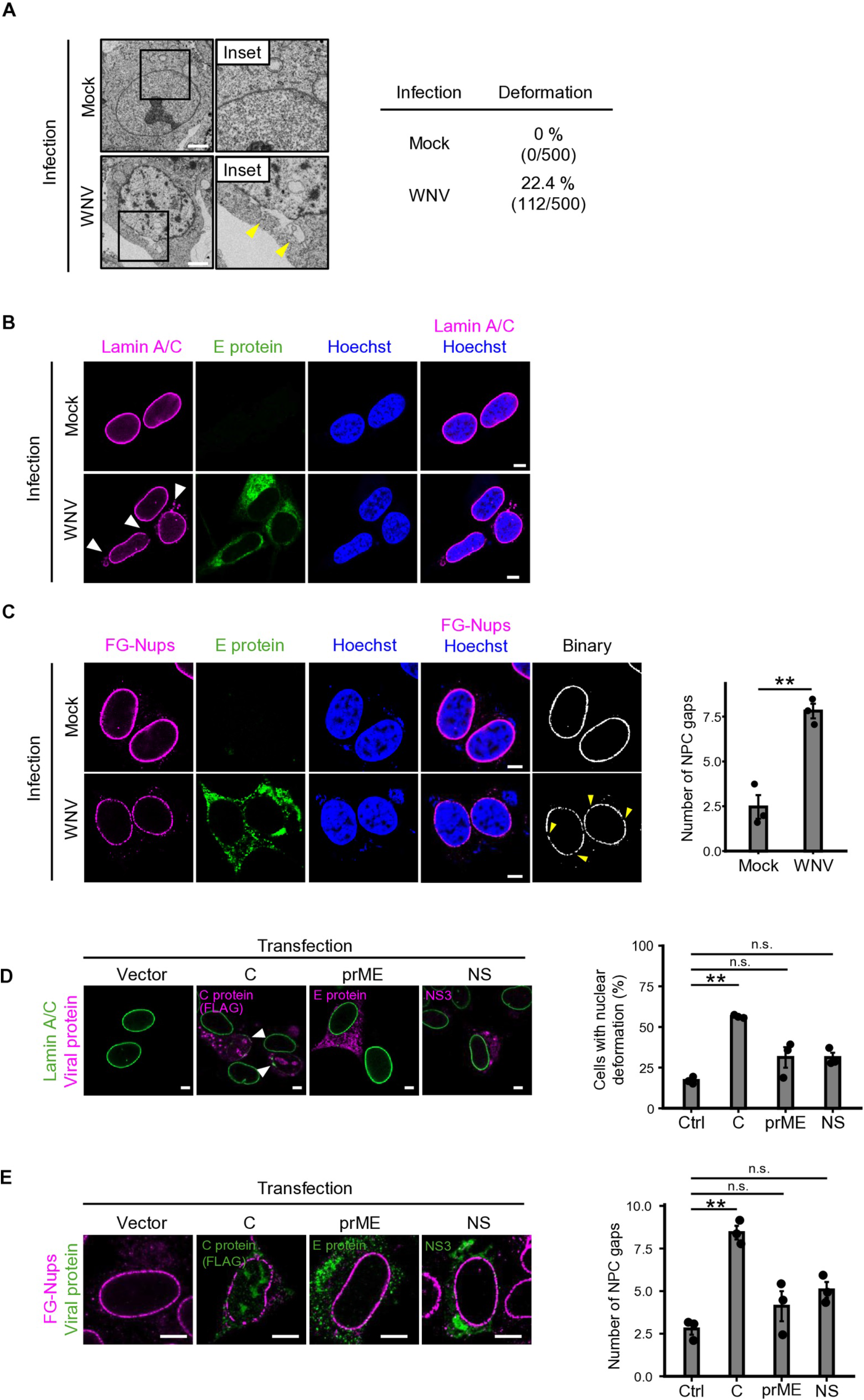
WNV infection induces nuclear deformation. (A) Transmission electron microscopy (TEM) analysis of SH-SY5Y cells infected with WNV. SH-SY5Y cells were infected with mock or WNV at 0.1 PFU/cell. The cells were fixed at 48 h post-infection (hpi) and subjected to TEM analysis. Arrowheads indicate discontinuities and bleb-like structures of the nuclear membrane (left). The ratio of cells with nuclear deformation was quantified (Mock: 0%, 0/500; WNV: 22.4%, 112/500) (right). Scale bar; 2 µm. (B) The lamin morphology in SH-SY5Y cells infected with WNV. SH-SY5Y cells were infected with WNV at 0.1 PFU/cell and the cells were fixed at 48 hpi and stained for E protein (green) and lamin A/C (magenta).

Arrowheads indicate the morphological changes of lamin A/C. Scale bar: 5 µm. (C) Distribution of the nuclear pore complex (NPC) in SH-SY5Y cells infected with WNV. SH-SY5Y cells were infected with WNV at 0.1 PFU/cell and the cells were fixed at 48 hpi and stained for E protein (green) and lamin A/C (magenta). Arrowheads indicate the regions lacking FG-Nups signals (NPC gaps). The number of NPC gaps per nucleus in binary images was quantified using Fiji image software. Binary images of FG-Nups staining were created by Otsu’s method using Fiji software. Scale bar: 5 µm. The bars indicate the mean ± standard error from three biological replicates. (D–E) The distribution of nuclear envelope components in SH-SY5Y cells expressing viral proteins. SH-SY5Y cells were transfected with the empty vector (Vector) or plasmids expressing One-STrEP-FLAG (OSF)-tagged C, prME, or all nonstructural (NS) proteins and harvested at 48 h post-transfection (hpt). The cells were stained for lamin A/C (D, green) or FG-Nups (E, magenta) with C (FLAG), E, or NS3 protein (D, magenta; E, green). Scale bar: 5 µm. Quantification was performed as described for (C). The bars indicate the mean ± standard error from three biological replicates. Statistical significance was evaluated using a two-tailed Welch’s t-test (C) or Dunnett’s test (**p < 0.01) (D and E).

The morphology of the nuclear membrane is mainly maintained by nuclear lamina, a filamentous meshwork of A- (lamin A/C) and B-type lamins (lamins B1 and B2) located beneath the nuclear inner membrane. The nuclear membrane is studded with nuclear pore complexes (NPCs) composed of approximately 30 nucleoporins (Nups) that regulate nucleocytoplasmic protein transport (NCT) [35]. To investigate the mechanisms underlying WNV-induced nuclear envelope deformation, we examined the localization of lamin A/C and Nups in WNV-infected cells. Lamin A/C formed a continuous and uniform ring surrounding the nucleus in mock-infected cells, whereas it exhibited a discontinuous pattern with accumulation near the nucleus in WNV-infected cells (Fig 1B). Similarly, the distribution of phenylalanine/glycine-repeat nucleoporins (FG-Nups) in mock-infected cells showed a continuous ring around the nucleus, whereas multiple gaps lacking FG-Nups signals were observed in WNV-infected cells (Fig 1C). The number of gaps identified using binary images in WNV-infected cells was higher than that in mock-infected cells (Fig 1C). In addition to FG-Nups, WNV-infected cells exhibited gap formation in Nup107 or POM121, which are scaffold and transmembrane Nups, respectively (S1A and B Fig). These results suggest that WNV infection induces nuclear deformation.

Nuclear deformation leads to NCT dysfunction [29]. Ras-related nuclear protein (Ran), a key regulator of NCT, is typically concentrated in the nucleus to support NCT. However, mislocalization of Ran to the cytoplasm results in the impairment of NCT [36]. Indeed, Ran was predominantly localized to the nucleus of mock-infected cells, whereas it was detected in both the nucleus and cytoplasm in WNV-infected cells (S2A Fig). The cytoplasmic-to-nuclear (C/N) fluorescence ratio of Ran in WNV-infected cells was significantly higher than that in mock-infected cells (S2A Fig). This indicates that WNV infection disrupted the nucleocytoplasmic distribution of Ran. Consistent with these findings, a GFP reporter containing both nuclear export and import signals (NES–GFP–3NLS), which alternates between the cytoplasm and nucleus, was detected in the cytoplasm of WNV-infected cells (S2B Fig). These results suggested that WNV infection disrupts NCT function.

Next, we sought to identify the viral protein responsible for nuclear deformation. The distribution of lamin A/C and FG-Nups was analyzed in cells expressing C, prME, or nonstructural (NS) proteins. The ratio of cells with lamin A/C morphological alterations and the number of FG-Nups gaps were significantly higher in C protein-expressing cells than in those expressing prME or NS proteins (Fig 1D and 1E). Similarly, the expression of C proteins derived from JEV, another orthoflavivirus, induced morphological alterations in lamin A/C and the formation of gaps in FG-Nups. In contrast, the C protein of TBEV only had minimal effects (S3A–D Fig). These results indicate that the WNV C protein is responsible for nuclear deformation, and that this property is shared among some orthoflaviviruses.

### WNV C protein induced phosphorylation and disassembly of nuclear lamina

Polymerization of A- and B-type lamins maintains nuclear lamina organization and nuclear morphology, whereas phosphorylation of lamins promotes their depolymerization [37]. To investigate the mechanisms underlying nuclear deformation induced by the C protein, we first examined the status of lamin polymerization based on the spatial relationship and interaction between lamin A/C and lamin B1. In cells transfected with the empty vector, lamin A/C and lamin B1 were colocalized to the nuclear periphery as a continuous ring, whereas the loss of both lamin A/C and lamin B1 was partially detected in C protein-expressing cells (Fig 2A). Next, the interaction between lamin A/C and lamin B1 was examined using co-immunoprecipitation. Lamin B1 was precipitated with an anti-lamin B1 antibody and the associated lamin A/C was detected with an anti-lamin A/C antibody. The amount of lamin A/C co-precipitated with lamin B1 was reduced in cells expressing the C protein compared to that in control cells (Fig 2B).

**Fig 2.**
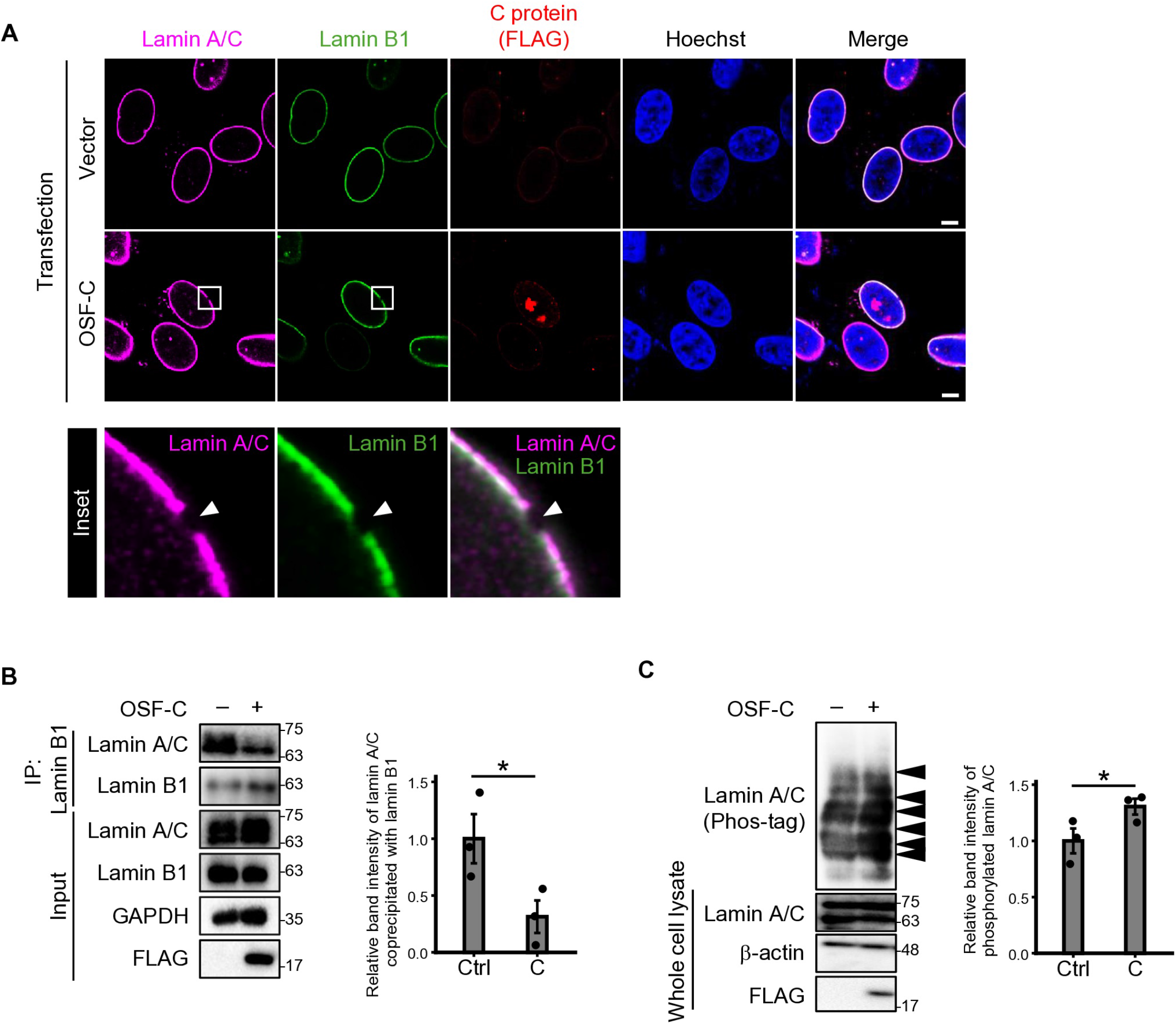
C protein induces depolymerization and phosphorylation of lamin proteins. (A) The spatial relationship between lamin A/C and lamin B1. SH-SY5Y cells were transfected with an empty plasmid (Vector) or a plasmid expressing OSF-C and fixed at 48 hpt. Cells were stained for lamin A/C (magenta), lamin B1 (green), and C protein (FLAG) (red). The inset highlights representative regions in which both lamins were depleted, indicated by the arrowhead. Scale bar: 5 μm. (B–C) SH-SY5Y cells expressing doxycycline-inducible OSF-C were treated with mock or 1 mg / mL doxycycline for 48 h and harvested. The cell lysate was subjected to anti-lamin B1 immunoprecipitation (B) and Phos-tag SDS–PAGE (C). (B) Co-immunoprecipitation analysis of lamin A/C and lamin B1. The whole-cell lysates were immunoprecipitated using anti-lamin B1 antibody, and the precipitated lamin A/C was analyzed by immunoblotting (left). Relative band intensity of immunoprecipitated lamin A/C was quantified (right). Each dot represents the mean value from three biological replicates and bars indicate the mean ± standard error. (C) Phosphorylation status of lamin A/C, as analyzed by Phos-tag SDS-PAGE. Arrowheads indicate bands corresponding to phosphorylated lamin A/C bands (left). Relative band intensity of phosphorylated lamin A/C was quantified (right). The bars indicate the mean ± standard error from three biological replicates. Statistical significance was evaluated using a two-tailed paired t-test (B and C) (*p < 0.05).

Next, we examined the phosphorylation status of lamin A/C in cells expressing the C protein using Phos-tag SDS-PAGE, which separates phosphorylated proteins according to their phosphorylated state [38]. The slow-migrating bands, which corresponded to phosphorylated lamin A/C [39], increased in cells expressing the C protein compared to that in control cells (Fig 2C). Taken together, these results suggest that the C protein enhances the phosphorylation of lamin and disassembly of the nuclear lamina.

### PKC α interacting with C protein was located in the perinuclear region

Because lamin phosphorylation is regulated by host kinases such as PKC, including PKCα, PKCδ, and PKCγ [40], we examined the subcellular localization of these PKC isoforms in cells expressing the C protein. Although PKCα was localized primarily around plasma membrane or cytoplasm in cells transfected with empty vector, PKCα in cells expressing the C protein was distributed in the cytoplasm and nucleus. Consistent with these observations, the amount of PKCα in the nucleus was significantly elevated in cells expressing the C protein (Fig 3B). Among other PKC isoforms, C protein expression did not affect the localization of PKCδ and PKCγ (S4 Fig). Furthermore, endogenous PKCα co-precipitated with the C protein (Fig 3C). We investigated the relationship between the subcellular interactions among the C protein, PKCα, and lamin A/C using a proximity ligation assay (PLA). PLA enables the detection of *in situ* protein–protein interactions by producing signals when two proteins are located within 40 nm of each other. In cells expressing the C protein, PLA signals between the C protein and PKCα, or lamin A/C were detected in the perinuclear region (Fig 3D and 3E). In addition, PLA signals between PKCα and lamin A/C were co-localized with the C protein in the perinuclear region (Fig 3F). The number of PLA signals was markedly increased in cells expressing the C protein compared to that in cells transfected with the empty vector (Fig 3G). These results suggested that the C protein interacted with PKCα on the nuclear lamina and promoted the proximity between PKCα and lamin A/C.

**Fig 3.**
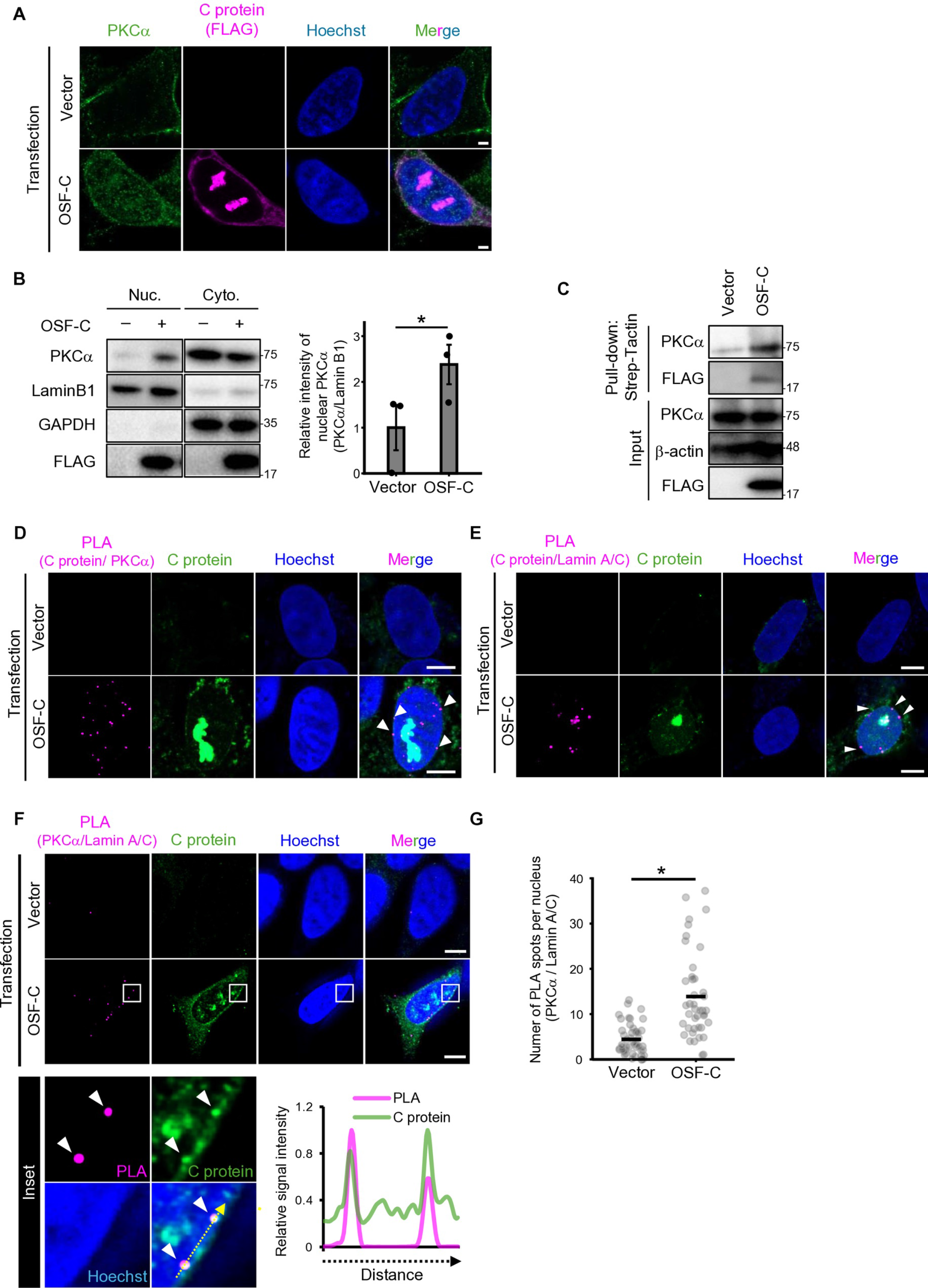
Protein kinase Cα interacting with the C protein was located in the perinuclear region. (A) Intracellular localization of PKCα. SH-SY5Y cells were transfected with an empty plasmid (Vector) or a plasmid expressing OSF-C and fixed at 48 hpt. Cells were stained for PKCα (green) and the C protein (FLAG) (magenta). Scale bar: 5 μm. (B) Subcellular fractionation of PKCα. SH-SY5Y cells transfected with an empty plasmid (Vector) or a plasmid expressing OSF-C were fractionated into nuclear and cytoplasmic fractions at 48 hpt and analyzed by immunoblotting (left). Relative intensity of PKCα normalized to that of lamin B1 was quantified (right). The bars indicate the mean ± standard error from three biological replicates. (C) Pull-down assay of interaction between the C protein and PKCα. HEK-293T cells were transfected with an empty plasmid (Vector) or a plasmid expressing OSF-C and harvested at 48 hpt. The cell lysates were subjected to Strep-Tactin pull-down assay, and the PKCα coprecipitated with OSF-C was detected by immunoblotting. (D–G) Subcellular interaction among C protein, PKCα, and lamin A/C. SH-SY5Y cells were transfected with an empty plasmid (Vector) or a plasmid expressing OSF-C and fixed at 48 hpt. The cells were subjected to the proximity ligation assay (PLA) using antibodies against PKCα and FLAG (C protein) (D), FLAG (C protein), and lamin A/C (E) or PKCα and lamin A/C (F). The OSF-C protein was stained with anti-C protein antibody. Scale bar: 5 µm. Arrowheads in the inset indicate colocalization of PLA signals with C protein.

Fluorescence intensity was measured along the yellow dotted arrow, and relative intensity was calculated by normalizing to the maximum fluorescence intensity. (G) The numbers of PLA spots between PKCα and lamin A/C per nucleus, as counted using Fiji image software. Each dot represents an individual cell from 40–50 cells. Data were analyzed using a two-tailed paired Student’s t-test (B) or Welch’s t-test (C and G) (*p < 0.05).

### PKC α mediated nuclear deformation induced by the C protein

We further investigated whether PKCα is required for C protein-induced nuclear deformation. First, the phosphorylation status of lamin A/C was examined in cells with downregulated PKCα expression. C protein expression increased the slow-migrating bands corresponding to phosphorylated lamin A/C in control siRNA-treated cells, whereas these bands were reduced in cells with downregulated PKCα expression (Fig 4A). Next, lamin morphology was examined in cells with downregulated PKCα expression. Morphological alterations induced by C protein expression were suppressed in cells exhibiting downregulated PKCα expression (Fig 4B). The number of cells with C protein-induced nuclear deformation was significantly reduced in cells with downregulated PKCα expression (Fig 4C). Furthermore, we examined whether the downregulation of PKCα expression rescued the NCT dysfunction induced by the C protein. Cytoplasmic mislocalization of NES–GFP–3NLS, which was observed in cells expressing the C protein with control siRNA, was rarely observed in cells with downregulated PKCα expression (S5A Fig). Accordingly, the cytoplasmic-to-nuclear fluorescence intensity ratio of GFP was significantly reduced in cells with downregulated PKCα expression compared to that in control siRNA-treated cells (S5B Fig).

**Fig 4.**
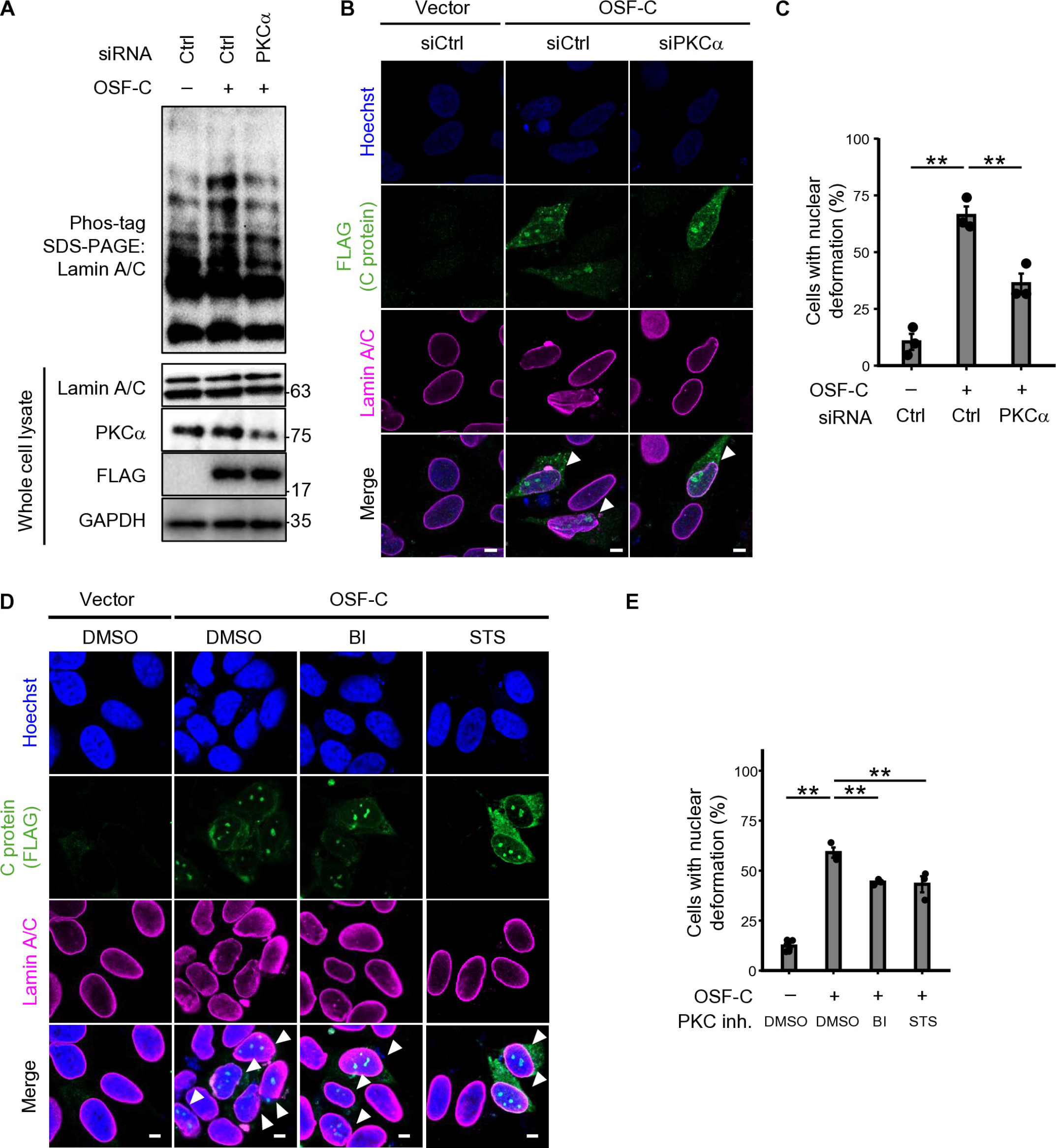
PKCα mediated lamin phosphorylation and nuclear deformation induced by C protein. (A) Phosphorylation status of lamin A/C. SH-SY5Y cells expressing doxycycline-inducible OSF-C were treated with control or PKCα siRNAs for 24h, and C protein expression was induced with doxycycline (1 μg/mL) for 48 h. Whole cell lysates were analyzed by Phos-tag SDS-PAGE and immunoblotting. (B) Effects of the downregulation of PKCα expression on nuclear lamina morphology. SH-SY5Y cells were transfected with an empty vector (Vector) or a plasmid expressing OSF-C 24 h after transfection with siRNAs. The cells were fixed at 48 hpt and stained for lamin A/C (magenta) and C protein (FLAG) (green). Arrowheads indicate cells expressing the C protein. Scale bar: 5 μm. (C) Quantification of the percentage of cells with deformed lamin A/C. The bars indicate the mean ± standard error from three biological replicates. (D) Effects of treatment with PKC inhibitors on nuclear morphology. SH-SY5Y cells transfected with an empty vector (Vector) or a plasmid expressing OSF-C and the culture medium was exchanged to medium containing PKC inhibitors, bisindolylmaleimide (BI) (1 mM) and staurosporine (STS) (10 nM) at 8 hpt. The cells were fixed at 32 hpt and stained for lamin A/C (magenta), C protein (FLAG) (green). Arrowheads indicate cells expressing the C protein. Scale bar: 5 μm. (E) Quantification of the percentage of cells with deformed lamin A/C. The bars indicate the mean ± standard error from three biological replicates. Statistical analysis was performed using Dunnett’s multiple comparison test compared to DMSO-treated cells expressing OSF-C (**p < 0.01) (C and E).

Next, we examined the effect of the pharmacological inhibition of PKC activity on nuclear morphology. C protein-mediated morphological alterations in lamin were inhibited in cells treated with the PKC inhibitors bisindolylmaleimide I (BI) or staurosporine (STS) (Fig 4D). The number of cells with nuclear deformation upon C protein expression was significantly reduced in cells treated with PKC inhibitors compared to that in DMSO-treated cells (Fig 4E). Taken together, these results suggest that PKCα is required for C protein-induced nuclear deformation.

### Inhibition of PKCα-dependent nuclear morphology attenuated viral multiplication in vitro

Because PKCα was related to nuclear deformation induced by the C protein, we subsequently investigated the importance of nuclear deformation in WNV multiplication. To this end, viral multiplication was examined in SH-SY5Y cells treated with siRNA against PKCα or PKC inhibitors. Viral titers were significantly lower in cells with downregulated PKCα expression than in control siRNA-treated cells (Fig 5A). Similarly, viral titers were significantly reduced in a dose-dependent manner in PKC inhibitor-treated cells compared to those in DMSO-treated cells (Fig 5B). The viability of cells with downregulated PKCα expression or those treated with PKC inhibitors was comparable to that of control siRNA- or DMSO-treated cells (S6A and B Fig). These results suggest that PKCα suppression inhibits WNV multiplication. Because nuclear deformation was observed in cells expressing the C proteins of WNV and JEV, but not TBEV (S2 Fig), we examined whether the inhibition of PKCα affected viral replication across other orthoflaviviruses. Treatment with BI significantly reduced the viral titers of JEV (S7A Fig) but had a minimal effect on TBEV (S7B Fig), suggesting that inhibition of viral replication by PKC inhibitors correlated with the presence of nuclear deformation phenotypes.

**Fig 5.**
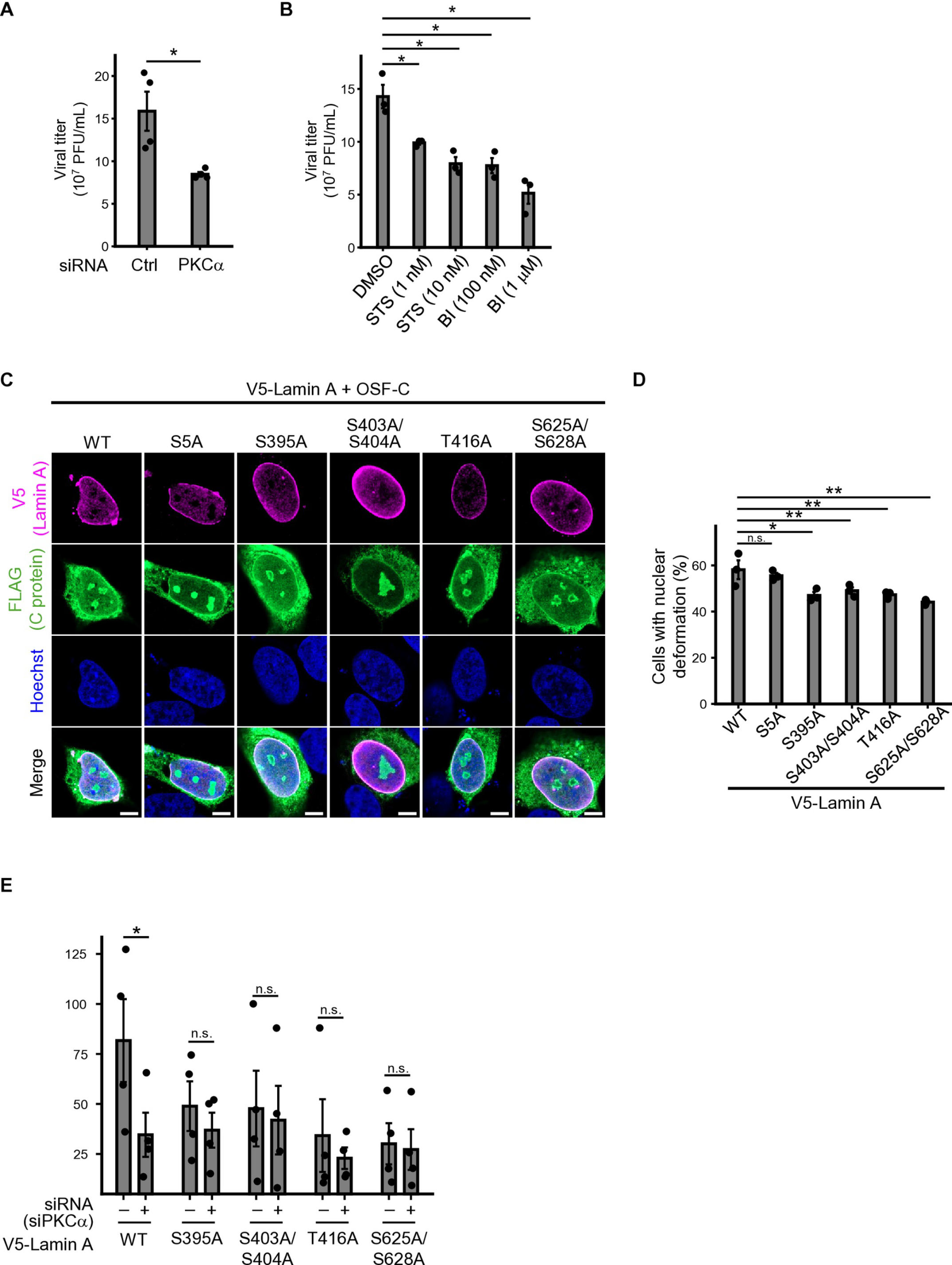
**Inhibition of nuclear deformation mediated by PKC**α **was involved in WNV replication.** (A) The effect of the downregulation of PKCα expression on WNV multiplication. SH-SY5Y cells were transfected with control or PKCα siRNAs for 24 h and infected with WNV at 0.1 PFU/cell. Culture supernatants were collected at 48 hpi and viral titers were measured by plaque assay. The bars indicate the mean ± standard error from four biological replicates. (B) The effect of PKC inhibition on WNV multiplication. SH-SY5Y cells were pre-treated with staurosporine (STS; 1 or 10 nM) or bisindolylmaleimide I (BI; 100 nM or 1 μM) for 1 h prior to infection with WNV (0.1 PFU/cell). The inhibitors remained in the medium during infection until 48 hpi. The viral titers in the supernatants were determined by plaque assay. The bars indicate the mean ± standard error from three biological replicates. (C) The effect of the C protein on nuclear morphology in cells expressing lamin A carrying serine- or threonine-to-alanine substitutions. SH-SY5Y cells were transfected with a plasmid expressing OSF-C and a plasmid expressing V5-tagged wild-type lamin A or lamin A mutants. The cells were stained for V5 (Lamin A) (magenta) and FLAG (C protein) (green). Scale bar: 5 μm (D) Quantification of the percentage of nuclear deformation in cells expressing wild-type or serine- or threonine-to-alanine substituted lamin A mutants. The bars indicate the mean ± standard error from three biological replicates. (E) Viral replication in cells expressing lamin A carrying serine- or threonine-to-alanine substitutions. SH-SY5Y cells stably expressing V5-tagged wild type or lamin A mutants (S395A, S403/S404A, T416A, or S625A/S628A) were infected with WNV at 0.1 PFU/cell 24h after transfection with control or PKCα siRNAs. Culture supernatants were collected at 48 hpi and viral titers were measured by plaque assay. The bars indicate the mean ± standard error from four biological replicates. Statistical analysis was performed using a two-tailed paired t-test (A and E) or Dunnett’s multiple comparison test (**p < 0.01, *p < 0.05, n.s., not significant) (B and D).

Next, we used lamin A mutants that were resistant to C protein-induced nuclear deformation to assess whether the PKCα-mediated enhancement of WNV proliferation depends on nuclear deformation through lamin phosphorylation. Several serine or threonine residues in lamin A regulate lamin organization through phosphorylation. Some of these residues are implicated in PKC-mediated lamin phosphorylation [17,41,42]. C protein-induced nuclear deformation was suppressed in cells expressing lamin A mutants carrying alanine substitutions at S395, S403 and S404, T416, or S625 and S628, but not in cells expressing the S5A mutant (Fig 5C and D). Subsequently, we analyzed WNV multiplication in cells expressing these nuclear deformation-resistant lamin A mutants. Viral titers were reduced in cells expressing the nuclear deformation-resistant lamin A mutants compared to those in cells expressing wild-type lamin A (Fig 5E). Furthermore, the downregulation of PKCα expression reduced the viral titer in cells expressing wild-type lamin A, whereas no marked reduction in viral titer was observed following the downregulation of PKCα expression in cells expressing nuclear deformation-resistant lamin A mutants (Fig 5E). These results suggest that C protein-induced PKCα-dependent nuclear deformation promotes WNV multiplication.

### Inhibition of PKCα-dependent nuclear deformation by PKC inhibitors suppressed disease progression of WNV infection in vivo

We examined the effects of PKCα-dependent nuclear deformation on WNV pathogenesis in mice. C57BL/6JJ mice were intraperitoneally inoculated with WNV and intracranially treated with BI at 3 and 6 days post-inoculation. Survival rate was significantly higher in mice treated with BI than in DMSO-treated mice (Fig 6A). No significant differences in viral titers were observed in the brain or spleen at 5 days post-inoculation between the two groups, whereas viral titers in the brain were significantly lower in BI-treated mice than in DMSO-treated mice at 7 days post-inoculation (Fig 6B). In the brains of infected mice at 7 days post inoculation, nuclear invaginations and wrinkles indicating nuclear membrane disruption [43] were observed in WNV-infected cells (Fig 6C). The number of cells with nuclear deformation in the brains of BI-treated mice was significantly lower than that in DMSO-treated mice (Fig 6D). These results suggested that PKCα-dependent nuclear deformation is associated with the neuropathogenesis of WNV infection *in vivo*.

**Fig 6.**
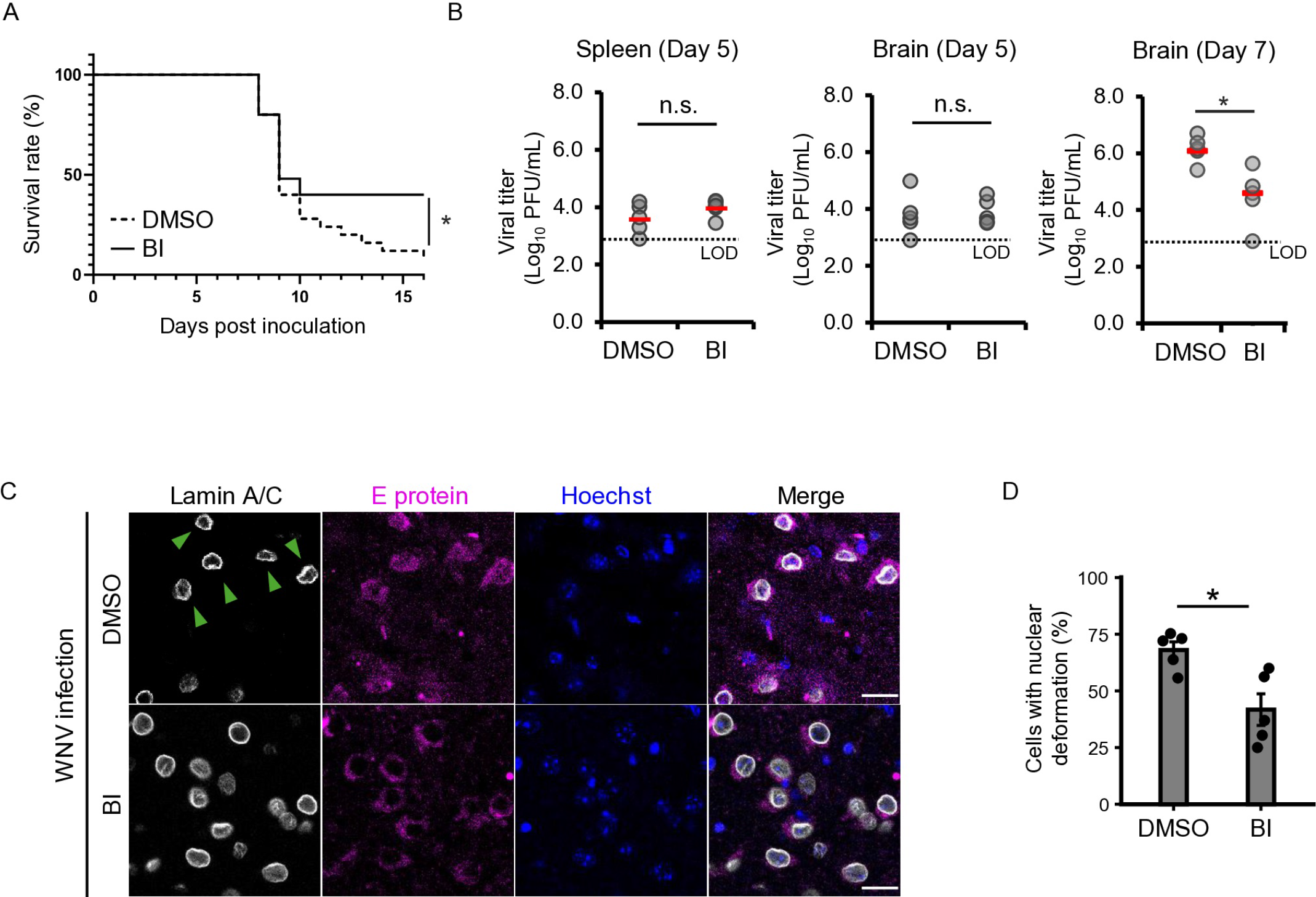
Inhibition of PKCα-dependent nuclear deformation by PKC inhibitors suppressed WNV disease progression *in vivo*. (A–D) Five-week-old female C57BL/6JJ mice were intraperitoneally inoculated with 100 pfu of WNV and intracranially administered DMSO or bisindolylmaleimide I (BI) at 3 and 6 days post infection (dpi). (A) The Kaplan–Meier survival curves (DMSO; n = 25, BI; n = 25). Survival differences were analyzed using a log-rank test (*p < 0.05). (B) Viral titers, as measured at 5 and 7 dpi in the brain and at 5 dpi in the spleen (DMSO; n = 5, BI; n = 5). Red bars indicate mean values; LOD, limit of detection. (C) Nuclear deformation in the brain. Sections of the cerebral cortex of WNV-inoculated mice treated with DMSO or BI were stained for lamin A/C (green) and the WNV antigen (magenta). Nuclei were counterstained with Hoechst. Arrowheads indicate abnormal lamin A/C morphology. (D) Quantification of the number of cells with nuclear deformation indicating nuclear invaginations or wrinkles. Each dot represents the mean value obtained from an individual mouse, and bars indicate the mean ± standard error for each group. Data were analyzed using a two-tailed Welch’s t-test (*p < 0.05) (B and D).

## Discussion

The C protein of orthoflavivirus is localized to the nucleus and promotes viral replication [7–11]. However, its effects on the nucleus and the mechanisms underlying its contribution to viral replication remain unclear. In this study, we showed that WNV C protein induces nuclear deformation. This deformation was driven by nuclear lamina disassembly following PKCα-mediated phosphorylation, which was triggered by interaction between PKCα and the C protein. Furthermore, the replication and pathogenesis of WNV were attenuated through suppression of nuclear deformation. These findings highlight the importance of nuclear deformation mediated by viral–host protein interactions during orthoflavivirus infection.

The C protein interacted with PKCα and increased the amount of this protein in nucleus. In addition, the C protein, PKCα, and lamin were localized near the nuclear envelope. Proteins of the PKC family, including PKCα, are host kinases that phosphorylate lamin proteins [44,45]. Phosphorylation of lamin causes disassembly of the polymerized lamin, resulting in its depletion and aggregation at the nuclear envelope [46], as observed in our results. Disassembly of lamins reduces the mechanical integrity of the nuclear envelope, leading to nuclear deformation and rupture [47,48]. PKCα alternates between the cytoplasm and the plasma membrane [49]. In DNA virus infections, such as herpes simplex virus 1 and cytomegalovirus, PKC is recruited to the perinuclear region from the cytoplasm through interactions with viral proteins, leading to lamin phosphorylation [21,50]. The WNV C protein alters the localization of host proteins to other cellular components through its interaction [51]. Thus, C protein may interact with PKCα in the cytoplasm or at the plasma membrane and subsequently translocate to the nucleus together with PKCα. However, PKCα is also localized to the nucleus through its activation [52]. This suggests that PKCα is first translocated into the nucleus by activation and is subsequently retained in the perinuclear region through interactions with the C protein. This accumulation of PKCα in the nucleus can promote phosphorylation of the nuclear lamina and nuclear deformation. Notably, nuclear deformation was induced by both the WNV C protein and the JEV C protein. Given that the WNV C protein has approximately 66.7% amino acid sequence identity with the JEV C protein and shares a similar subcellular localization [53,54], the shared elements of these orthoflaviviruses may be involved in PKCα translocation and nuclear deformation. However, additional analysis of the spatiotemporal relationship between PKCα and the C protein of orthoflaviviruses is required to further elucidate the mechanisms underlying WNV-induced nuclear deformation.

Nuclear deformation potentially facilitates viral multiplication via two mechanisms. The first is that WNV-induced nuclear deformation alters cellular gene expression. Lamins interact with chromatin at the nuclear periphery and regulate chromatin organization and transcriptional activity [55]. In neuronal cells, structural alterations in lamins or invagination of the nuclear envelope led to genomic DNA breaks, heterochromatin relaxation, and alterations in gene expression [30,56,57]. Furthermore, mutations in lamin A are associated with progeria and cardiomyopathy, known as laminopathy. These mutations alter genome organization, histone methylation, and gene expression [58,59]. Moreover, phosphorylated lamin alters the binding region of chromatin and upregulates the expression of genes that define the cellular phenotype [60]. We showed that the C protein of JEV also induced nuclear deformation, and the C protein was reported to influence cellular RNA expression profiles [61]. These findings suggest that orthoflaviviruses, including WNV, promote their multiplication by regulating host gene expression through nuclear deformation. The second potential mechanism involves the mislocalization of proteins as a consequence of the disruption of nucleocytoplasmic protein transport. Type I interferon (IFN), a central component of the innate immune response, is produced by the translocation of transcription factors such as IFN regulatory factor 3 (IRF3) into the nucleus and binding to IFN promoters. Many viruses, such as picornaviruses, SARS-CoV-2, and porcine reproductive and respiratory syndrome virus (PRRSV), suppress IFN expression by impairing nucleocytoplasmic transport through the disruption of NPC integrity and nuclear-to-cytoplasmic Ran gradient [62–65]. In addition, nuclear proteins such as DDX56, a protein associated with DEAD-box RNA helicases, or TIAR and TIA1, stress response-related proteins, are distributed in the cytoplasm during WNV infection and promotes viral replication [66–68]. These observations suggest that WNV infection suppresses host antiviral responses and/or exploits cytoplasmic nuclear proteins by disrupting nucleocytoplasmic transport.

Suppression of PKCα expression or treatment with pharmacological PKC inhibitors attenuates the nuclear deformation caused by WNV infection and reduces viral replication and pathogenesis. PKCα is a multifunctional protein involved in diverse signaling pathways, including cell proliferation, differentiation, apoptosis, and inflammatory responses [69–72].

PKC inhibitors suppress intracellular WNV genome replication [73]. Furthermore, the inhibition affects the replication of other RNA viruses, such as influenza virus and PRRSV, by suppressing endocytic trafficking or inflammatory responses [74–76]. These findings suggest that the reduction in WNV replication upon PKC inhibition was mediated by mechanisms other than the suppression of nuclear deformation. Nevertheless, the expression of lamin A mutants, predicted to be resistant to PKC-mediated phosphorylation, attenuated WNV-induced nuclear deformation and viral replication. Moreover, the downregulation of PKCα expression did not further reduce WNV replication in cells expressing lamin mutants. These findings suggest that PKCα-mediated lamin phosphorylation contributes, at least in part, to C protein-induced nuclear deformation and the enhancement of WNV replication.

In summary, our findings provide the first evidence of the importance of nuclear deformation in orthoflavivirus infection and reveal the molecular mechanism underlying this process, which is mediated by the interaction between the C protein and PKCα. These findings offer a new perspective for the study of many other RNA viruses in which the nucleus is considered irrelevant to viral replication and pathogenesis.

## Materials and Methods

### Cells and viruses

SH-SY5Y cells derived from human neuroblastomas were obtained from the European Collection of Authenticated Cell Cultures and cultured in Dulbecco’s modified Eagle’s medium (DMEM): Nutrient Mixture F-12 (FUJIFILM Wako Pure Chemical Corporation, Osaka, Japan) supplemented with 10% heat-inactivated fetal bovine serum (FBS). HEK-293T cells derived from human embryonic kidney were obtained from Dr. Matsuura (Osaka University, Japan) and grown in high-glucose DMEM (FUJIFILM Wako Pure Chemical Corporation) supplemented with 10% heat-inactivated FBS. Vero cells derived from African green monkey kidneys were obtained from JCRB Cell Bank (Osaka, Japan) and cultured in Eagle’s minimum essential medium (EMEM) (FUJIFILM Wako Pure Chemical Corporation) supplemented with 10% heat-inactivated FBS. Baby hamster kidney (BHK) cells were cultured in EMEM (FUJIFILM Wako Pure Chemical Corporation) supplemented with 10% heat-inactivated FBS.

The WNV 6-LP strain (accession number: AB185914.2) was established from the WNV NY99-6922 strain isolated from mosquitoes in 1999, as described previously [77]. Vero cells were inoculated with WNV and the supernatants were harvested 3 days post-infection (dpi). The JEV JaGAr-01 and TBEV Oshima 5-10 strains (accession numbers: AF069076.1, AB062063.2, respectively) were propagated in BHK cells. Supernatants containing the virus were stored at -80°C until use. All viral experiments were performed at a biosafety level-3 (BSL-3) facility at Hokkaido University, Japan, in accordance with institutional guidelines.

### Plasmid construction

A plasmid encoding the WNV C protein with an N-terminal One-STrEP-FLAG tag (pCMV-OSF-C) was previously constructed [78]. Plasmids expressing prME (pCXSN-prME) and the subgenomic RNA of the WNV (pCMV-WNrep) were also constructed previously [79]. The PKCα sequence was amplified from the total RNA of SH-SY5Y cells using a Prime Script One Step RT-PCR Kit Ver.2 (Takara Bio Inc., Shiga, Japan). The lamin A sequence was amplified using the KOD One PCR Master Mix (Toyobo, Osaka, Japan) from the cDNA synthesized using the total RNA of SH-SY5Y cells with PrimeScript II 1st strand cDNA Synthesis Kit (Takara Bio Inc.). Amplified PKCα and lamin A sequences were introduced between the XhoI and NotI sites downstream of the V5 tag in the pCMV-V5 plasmid using the In-Fusion HD cloning kit (Takara Bio Inc.). The resultant constructs were designated as pCMV-V5-PKCα or pCMV-V5-lamin A. Variants of the plasmid expressing serine or threonine-to-alanine substitution mutants of lamin A protein were generated via inverse PCR using appropriate primers. The lentiviral packaging plasmids pCAG-HIVgp and pCMV-VSV-G-RSV-Rev and the lentiviral vector CSII-CMV-MCS-IRES2-Bsd were kindly provided by Dr. Hiroyuki Miyoshi (Riken, Tsukuba, Japan). The coding sequence of V5-tagged lamin A was amplified from pCMV-V5-lamin A using the KOD One PCR Master Mix (Toyobo) and inserted into the CSII-CMV-MCS-IRES2-Bsd vector using an In-Fusion HD cloning kit (Takara Bio Inc.). The resultant construct was designated as CSII-CMV-V5-lamin A-IRES2-Bsd.

### Antibodies, reagents, and transfection

Rabbit anti-JEV serum exhibiting cross-reactivity with the WNV E protein was produced as previously described [80]. Rabbit anti-JEV C protein that exhibited cross-reactivity with the WNV C protein was kindly provided by Dr. Morita (Hirosaki University, Japan). Rabbit anti-NS3 serum was prepared as previously described [81]. The following antibodies were purchased: rabbit anti-Lamin B1 polyclonal antibody (MBL, Nagoya, Japan); mouse anti-Nuclear pore complex (Mab414) monoclonal antibody (BioLegend, CA, USA); mouse anti-Lamin A/C (4C11) monoclonal antibody (Cell Signaling Technology, MA, USA); rabbit anti-POM121 polyclonal antibody (Proteintech, IL, USA); rabbit anti-Nup107 polyclonal antibody (Proteintech); rabbit anti-Ran polyclonal antibody (Proteintech); rabbit anti-DDDDK-tag polyclonal antibody (MBL or Proteintech); rabbit anti-PKCα polyclonal antibody (Proteintech); rabbit anti-PKCɣ polyclonal antibody (Proteintech); rabbit anti-PKCδ polyclonal antibody (Proteintech); mouse anti-Beta actin monoclonal antibody conjugated with HRP (Proteintech); and mouse anti-GAPDH monoclonal antibody conjugated with HRP-DirectT (MBL).

Plasmids were transfected into HEK-293T or SH-SY5Y cells using the X-tremeGENE HP DNA Transfection Reagent (Roche, Mannheim, Germany) or Lipofectamine 3000 Transfection Reagent (Invitrogen) according to the manufacturers’ instructions. siRNA were transfected with SH-SY5Y using the Lipofectamine RNAiMax Transfection Reagent (Invitrogen).

### Generation of doxycycline-inducible OSF-C-expressing SH-SY5Y cells

Doxycycline-inducible OSF-C-expressing cells were generated using the Retro-X™ Tet-One™ Inducible Expression System (Takara Bio Inc.) according to the manufacturer’s instructions. The coding sequence of the WNV C protein, fused to an N-terminal OSF tag, was amplified by PCR and cloned into the pRetroX-Tet-One vector (pRetroX-Tet-One-OSF-C) using an In-Fusion HD cloning kit (Takara Bio). Retroviral particles were produced by transfecting GP2-293 packaging cells with the pRetroX-Tet-One-OSF-C plasmid using Lipofectamine 3000 (Thermo Fisher Scientific). At 48 h post-transfection, virus-containing supernatants were collected and clarified by centrifugation at 500 × *g* for 3 min and applied to SH-SY5Y cells in the presence of 8 µg/mL polybrene (Sigma-Aldrich). Stable polyclonal populations were obtained by selection with puromycin (5 µg/mL) for 10–14 days. To induce OSF-C expression, doxycycline was added to the culture medium at a final concentration of 1 µg/mL for 48 h.

### TEM

SH-SY5Y cells were inoculated with WNV at a multiplicity of infection (MOI) of 1 and cultured for 48 h. The cells were scraped off the plate, pelleted by centrifugation at 3,000 rpm for 5 min, and fixed for 30 min with 2.5% glutaraldehyde and 2% paraformaldehyde (PFA) in 0.1 M phosphate buffer. The samples were then post-fixed in an osmium fixation solution [1% (w/v) osmium tetroxide, 1.25% (w/v) potassium ferrocyanide, and 5 mM calcium chloride]. After staining the blocks with 0.5% aqueous uranyl acetate, specimens were dehydrated and embedded in Spurr’s resin. Thin sections were mounted on copper grids and post-stained with saturated uranyl acetate and lead citrate solutions. The specimens were observed using an HT7700 transmission electron microscope (Hitachi High Technologies).

### Immunocytochemistry

Cells were fixed in 4% PFA or ice-cold ethanol at -20°C for 10 min and washed with phosphate buffered saline (PBS). Permeabilization and blocking were performed using 0.1% Triton X-100 and 1% bovine serum albumin (BSA) in PBS for 30 min at room temperature. Next, cells were incubated overnight at 4°C with the indicated primary antibodies diluted in 1% BSA–PBS, followed by incubation with Alexa Fluor 488-, 555-, or 647-conjugated secondary antibodies (Thermo Fisher Scientific, Waltham, MA, USA) for 1 h at room temperature. Zenon rabbit IgG labeling kits (Thermo Fisher Scientific) were used for the fluorescent labeling of rabbit primary antibodies. The nuclei were counterstained with Hoechst 33342. Images were acquired using an LSM800 confocal laser-scanning microscope (Carl Zeiss, Jena, Germany) with ZEN software (Carl Zeiss) or an APX100 fluorescence microscope with CellSens imaging software (Evident Scientific, Tokyo, Japan).

### Generation of SH-SY5Y cells stably expressing NES–GFP–3NLS or lamin A

SH-SY5Y cells stably expressing NES–GFP–3NLS or V5-tagged lamin A were generated using the lentiviral vectors. For lentiviral production, HEK-293T cells were transfected with pCAG-HIV-gp, pCMV-VSV-G-RSV-Rev, and CSII-CMV-NES-GFP-3NLS-IRES2-Bsd or CSII-CMV-V5-lamin A-IRES2-Bsd. At 48 h post-transfection, the culture supernatants were collected and added to SH-SY5Y cells. Stable polyclonal populations were obtained by selection with blastcidin (10 µg/mL) for 10–14 days.

### Immunoblotting

The cells were washed with PBS and lysed in lysis buffer (1% Triton X-100, 50 mM Tris-HCl [pH 7.5], 1 mM EDTA, and 0.25 M sucrose) containing the protease inhibitor (Nacalai Tesque, Kyoto, Japan). The lysates were centrifuged at 17,700 × *g* for 15 min at 4°C, and the resulting supernatants were mixed with an equal volume of 2× sample buffer (3% SDS, 10% glycerol, and 100 mM Tris-HCl [pH 6.8]). Total cell lysates were prepared in 1× sample buffer. The samples were separated by SDS-PAGE. Next, the proteins were transferred to a PVDF membrane, blocked with 5% skim milk in TBS-T (1% Triton, 25 mM Tris-HCl [pH 7.4], 0.14 M NaCl, and 2.7 mM KCl), and incubated overnight with each antibody at 4°C. After washing, membranes were incubated with horseradish peroxidase-conjugated secondary antibodies. Bands were visualized with a chemiluminescent HRP substrate (Millipore, Billerica, MA, USA) using a ChemiDoc XRS+ Imager (Bio-Rad, Hercules, CA, USA). Acquired images were analyzed with the Image Lab Software (Bio-Rad) and Fiji software.

### Protein Precipitation

SH-SY5Y cells transfected with pCMV-OSF-C were washed with PBS and lysed in lysis buffer. The lysates were centrifuged at 17,700 × *g* for 15 min at 4°C. The supernatants were rotated with Strep–Tactin Sepharose 4Flow high capacity (IBA Lifesciences, Göttingen, Germany) at 4°C for 2 h. Sepharose was washed with Buffer W (150 mM NaCl, 1 mM EDTA, and 100 mM Tris-HCl [pH 8.0]) and the bound proteins were eluted with 1× sample buffer and separated by SDS-PAGE. Lamin complexes were co-immunoprecipitated using the Capturem™ IP & Co-IP Kit (Takara Bio Inc.) according to the manufacturer’s instructions.

SH-SY5Y cells expressing doxycycline-inducible OSF-C were treated with 1 µg/mL doxycycline for 48 h or left untreated as a control. The cells were washed with ice-cold PBS and lysed in lysis buffer containing protease inhibitors on ice for 15 min. Lysates were clarified by centrifugation at 17,700 × *g* for 15 min at 4 °C. The supernatant was incubated with an anti-lamin B1 antibody for 20 min at room temperature. For immunoprecipitation, the anti-lamin B1 antibody was first immobilized on a Capturem IP column, and the column was washed several times with wash buffer. Bound proteins were eluted using the kit elution buffer and mixed with 2× sample buffer. The eluates were analyzed by SDS–SDS-PAGE and immunoblotting. Co-immunoprecipitation of lamin A/C with lamin B1 was detected using an anti-lamin A/C antibody.

### Phos-tag SDS-PAGE analysis

Phosphorylation of lamin A/C was analyzed by Mn²⁺–Phos-tag SDS–PAGE (FUJIFILM Wako Pure Chemical Corporation). Cell lysates were prepared in 1× sample buffer. The samples were separated on 5% SDS–polyacrylamide gels containing 20 µM Phos-tag acrylamide and 20 µM MnCl₂ (FUJIFILM Wako Pure Chemical Corporation). Following electrophoresis, gels were soaked three times in blotting buffer containing 10 mM EDTA for 10 min to remove Mn²⁺. Next, they were equilibrated in blotting buffer without EDTA for 10 min. Proteins were transferred to PVDF membranes and analyzed by immunoblotting using an anti-lamin A/C antibody (Cell Signaling Technology). Slower-migrating lamin A/C bands were interpreted as phosphorylated species, as previously described [39].

### PLA

PLA was performed using the Duolink *In Situ* Red Kit Mouse/Rabbit (Sigma-Aldrich), according to the manufacturer’s instructions. Cells were fixed with 4% PFA for 10 min at room temperature and permeabilized with 0.1% Triton X-100 in PBS. After blocking with Duolink blocking buffer for 1 h at 37°C, the cells were incubated overnight at 4°C with primary antibodies diluted in Duolink antibody dilution buffer. PLA PLUS and MINUS probes were applied for 1 h at 37°C, and ligation was performed with Ligation buffer for 30 min at 37°C. Signal amplification was then performed for 100 min at 37°C using polymerase diluted in Amplification buffer. To detect the C protein, cells were incubated overnight at 4°C with rabbit anti-JEV C protein serum cross-reactive with WNV C protein, which was labeled with Alexa Fluor 488 using a Zenon Rabbit IgG Labeling Kit (Thermo Fisher Scientific). Nuclei were counterstained with Hoechst 33342, and images were acquired using a Zeiss LSM 800 confocal microscope (Carl Zeiss).

### Viral titration

Subconfluent SH-SY5Y cells or SH-SY5Y cells stably expressing V5-lamin A were inoculated with WNV at an MOI of 0.1 at 24 h after transfection with siRNAs. For inhibitor treatment, SH-SY5Y cells were pre-incubated with DMSO or the PKC inhibitors BI (Selleckchem, TX, USA) or STS (Selleckchem) for 1 h and inoculated with WNV or JEV at an MOI of 0.1 or TBEV at an MOI) of 0.01. The inhibitors were maintained in the culture medium throughout the infection. Supernatants from infected cells were harvested 48 h post-infection (hpi) and stored at -80°C until use. For viral titration, the collected supernatants of cells inoculated with WNV, JEV, or TBEV were serially diluted and inoculated onto Vero cell monolayers for WNV and JEV titration or BHK cell monolayers for TBEV titration. After 1 h incubation at 37°C with rocking, the inoculum was removed. Next medium containing 5% FBS and 1.25% methyl cellulose or that containing 2% FBS and 1.5% carboxymethyl cellulose was added onto Vero or BHK cells, respectively. After 4 days of incubation, the plaques were visualized by staining with 0.25% crystal violet in 10% formalin.

### Cell viability assay

SH-SY5Y cells transfected with siRNA were seeded into 96-well plates and incubated for 48 h. For inhibitor treatment, SH-SY5Y cells were seeded into 96-well plates and incubated with 100 nM or 1 mM BI (Selleckchem) or 1 nM or 10 nM STS (Selleckchem) for 24 h. Cell viability was measured using the CellTiter-Glo Luminescent Cell Viability Assay (Promega Corporation, WI, USA) according to the manufacturer’s instructions. Briefly, CellTiter-Glo Reagent was added to the cells, followed by incubation for 3 min. Next, luminescence was measured using a GloMax Discover Microplate Reader (Promega Corporation).

### Ethics statement

All animal experiments were conducted in accordance with the basic guidelines for animal experiments of the Ministry of Education, Culture, Sports, Science, and Technology (MEXT), Japan. The President of Hokkaido University approved all animal experiments after review by the Institutional Animal Care and Use Committee of Hokkaido University (approval no. 23-0128).

### Inoculation of WNV into mice

Five-week-old C57BL/6JJmsSlc mice were purchased from Japan SLC, Inc. (Shizuoka, Japan). The animals were anesthetized with isoflurane and infected with WNV via intraperitoneal inoculation at 100 PFU/mouse. At 3 and 6 dpi, BI was intracranially administered at a dose of 4.12 µg/body in PBS, with a final DMSO concentration of 10%. Brains and spleens were collected after euthanasia at 5 or 7 dpi.

### Immunofluorescence staining of tissue samples

Collected tissue samples were fixed in 4% PFA and embedded in paraffin. The sections were deparaffinized using PathoClean (FUJIFILM Wako Pure Chemical Corporation) and a graded series of ethanol (99%, 95%, 90%, 80%, and 70%) was used for rehydration. Sections were subjected to antigen retrieval in 10 mM sodium citrate buffer (pH 6.0) in a pressure cooker. Subsequently, the sections were blocked with 10% normal goat serum for 10 min and incubated with primary antibodies diluted in Can Get Signal immunostain solution (Toyobo) at 4 °C overnight. The sections were then incubated with Alexa Fluor 488- or 555-conjugated secondary antibodies and Hoechst 33342 diluted in Can Get Signal immunostain solution (Toyobo) at room temperature for 1 h.

### Quantification and statistical analysis

Images acquired by immunocytochemistry were recorded using an APX100 fluorescence microscope (Evident Scientific) or LSM 800 confocal laser scanning microscope (Carl Zeiss). Images were processed using ZEN (Carl Zeiss), CellSens imaging (Evident Scientific), or Fiji software. To count the number of gaps in FG-Nups staining, the images were converted to binary images using Otsu’s thresholding method and processed with the closing operation in Fiji software. Nuclear or cytoplasmic regions were segmented using the cyto3 or nuclei model in Cellpose v3.1.0 [82]. Nuclear deformation was defined as nuclei showing abnormal morphology, including discontinuous lamin staining, irregular accumulation, or cytoplasmic dispersion of lamin. The fluorescence intensity in the segmented regions and number of cells with nuclear deformation were quantified using the Fiji software (https://github.com/fiji).

Immunofluorescence images of brain sections were recorded using an LSM 700 (Carl Zeiss) or APX100 fluorescence microscope (Evident Scientific). Nuclear deformation in brain sections was defined as a nuclei exhibiting wrinkles or invaginations, as described previously [43]. At least three fields were analyzed for each condition, and cell numbers were quantified using the Fiji software.

Densitometric quantification of the immunoblot bands for each lane was performed using the Image Lab software (Bio-Rad) or Fiji software, followed by normalization to the corresponding loading control from the same blot.

Data are presented as the mean ± standard error of the mean. Statistical analyses and data visualization were performed using R software. Comparisons between two groups were performed using a two-tailed Student’s or Welch’s t-test. Comparisons between multiple groups were performed using Dunnett’s test or the Holm-adjusted pairwise t-test. Survival curves were generated using the Kaplan–Meier method and compared using the log-rank test. Statistical significance was set at p < 0.05.

## Acknowledgments

We sincerely thank Dr. Hiroyuki Miyoshi for kindly providing CSII-CMV-MCS-IRES2-Bsd, pCAG-HIVgp, and pCMV-VSV-G-Rev, and all members of Sawa and Kariwa Laboratories for their valuable discussions. We also thank Ms. Sato for assistance with the experiments.

